# GABA and I_A_ Independently Regulate rNST Responses to Afferent Input

**DOI:** 10.1101/2021.09.14.460273

**Authors:** Z. Chen, D.H. Terman, S.P. Travers, J.B. Travers

## Abstract

Taste responses in the rostral nucleus of the solitary tract (rNST) influence motivated ingestive behavior via ascending pathways, and consummatory reflex behavior via local, brainstem connections. Modifications to the afferent signal within the rNST include changes in gain (the overall rate of neuron activity) and changes in gustatory tuning (the degree to which individual neurons respond to divergent gustatory qualities). These alterations of the sensory signal derive from both synaptic interactions within the nucleus and the constitutive cellular membrane properties of rNST neurons. GABA neurons are well represented within the rNST, as is expression of KV4.3, a channel for a rapidly inactivating outward K^+^ current (I_A_). GABAergic synapses suppress rNST responses to afferent input and previous studies showed that this suppression is greater in cells expressing I_A_, suggesting a possible interaction. Here, we examine the potential interaction between GABAergic inhibition and I_A_ channels in a series of patch clamp experiments. Optogenetic release of GABA suppressed rNST responses to afferent (electrical) stimulation and this effect was greater in cells with I_A_, confirming an earlier report. We further observed that the composite inhibitory postsynaptic potential was larger in I_A_ positive cells, suggesting one mechanism for the greater afferent suppression. Blocking I_A_ with the channel blocker AmmTX3, enhanced the response to afferent stimulation, suggesting a suppressive role for this channel in regulating afferent input at rest. However, pharmacologic blockade of I_A_ did not suppress GABAergic inhibition, indicating that I_A_ and GABA independently regulate excitatory afferent input.

## Introduction

Neurons in the rostral nucleus of the solitary tract (rNST) relay taste afferent input to the forebrain via a connection with the parabrachial nucleus and to local reflex pathways via projections to the caudal nucleus of the solitary tract (cNST) and subjacent reticular formation (1–5). Factors shaping the output signal from the rNST include convergence of primary afferents onto second order neurons (6–10), synaptic interactions between neurons within and outside the nucleus, and the intrinsic membrane properties of rNST neurons themselves (11, 12) reviewed in (13). GABA is one important modulator for this processing.

The rNST is endowed with an extensive GABAergic presence, arising from both local inhibitory interneurons and sources extrinsic to the nucleus (14–18). Indeed, immunohistochemical visualization of markers for GABA differentiates the nucleus from surrounding structures, (e.g. figure 3B (19)), emphasizing its importance. GABA_A_ receptors predominate in the rNST but there is a modest contribution of GABA_B_ receptors as well (20, 21). Activation of these receptors is implicated in gain control, providing inhibition that counteracts the purely excitatory afferent input (6, 10, 22). GABA inhibition may also play a role in gustatory tuning (23, 24).

In addition, many rNST projection neurons are endowed with a prominent transient outward potassium current (I_A_) (11, 12, 19, 25, 26). In the rNST I_A_ appears to be primarily mediated by KV4 channels and, similar to the distribution of GABA, immunohistochemical visualization for KV4.3 differentiates the nucleus from surrounding structures (e.g. figure 12A in (12)), again highlighting its potential importance. Although the presence of I_A_ in the rNST is well documented, its functional significance remains unclear. In the hippocampus, I_A_ channels located on distal dendrites suppress excitatory input (27) and in the cNST, cells with I_A_ are less responsive to afferent input compared to cells without (28). In the rNST, the response to afferent stimulation in non-GABAergic (putative projection) cells with I_A_ showed greater suppression in response to GABA activation suggesting a possible interaction between these two regulatory mechanisms (19). In theory, a feed-forward inhibitory circuit in which primary afferents excite GABAergic interneurons that in turn contact rNST projection neurons (19, 29), could make more Kv4.3 channels available so that the resulting I_A_ outward current is larger, amplifying the inhibitory effect of GABA. On the other hand, I_A_ could dampen excitatory input independently since I_A_ is operational at threshold, constituting a window current in NST cells (12, 30, 31)

In the present study we used whole cell patch clamp recording combined with optogenetics to identify non-GABAergic rNST neurons, a population which includes projection cells. We tested the hypothesis that there was an interaction between GABAergic inhibition and I_A_ by measuring the efficacy of inhibition in suppressing the response to afferent stimulation before and after blocking I_A_ with the specific KV4 channel blocker AmmTX3 (32, 33). We further implemented a dynamic clamp protocol to assess the impact of adding a synthetic I_A_ conductance on the response to afferent stimulation.

As expected, the optogenetic release of GABA suppressed afferent responses in non-GABA neurons and, confirming our previous observations, this effect was more pronounced in cells with I_A_ (19). We further observed that the degree of suppression in I_A_-positive cells (I_A+_) correlated with the magnitude of GABA-evoked inhibitory postsynaptic potentials (IPSPs). Manipulation of I_A_ channels also influenced the response to afferent stimulation. The pharmacological block of I_A_ enhanced the response to afferent stimulation whereas adding a synthetic I_A_ conductance with a dynamic clamp protocol reduced the response. Evidence for an interaction between GABAergic inhibition and I_A_, however, was lacking. Blocking I_A_ with AmmTX3 did not alter GABA inhibition, leading us to conclude that GABA inhibition and I_A_-mediated suppression of afferent input function independently.

## Methods

Experiments were conducted in mice (4-12 weeks: N= 70) that expressed channelrhodopsin (ChR2 H134R) in GABAergic neurons by placing the expression of ChR2 under the control of the promoter for either glutamic acid decarboxylase-65 (GAD65, also known as GAD2, n= 33 male, 32 female) or the vesicular GABA transporter, VGAT (n=2 male, 3 female). Animals were bred by crossing a Cre-dependent ChR2-EYFP mouse line (JAX #012569) with mice expressing Cre recombinase under the control of the endogenous promotor for GAD65 (GAD2-IRES-Cre, Jax 010802) or VGAT (VGAT-IRES-Cre, Jax 016962). Mice received ad libitum water and food until the day of the experiment. Experimental protocols were approved by the Ohio State University Institutional Animal Care and Use Committee in accordance with guidelines from the National Institutes of Health.

### Slice preparation

Following anesthetization with isoflurane, decapitation, and rapid removal and cooling of the brain, acute slices were prepared for electrophysiological recording. The brainstem was blocked, fixed to a ceramic block using cyanoacrylate glue, and coronal brainstem slices 250 μm thick were cut with a vibratome (model 1000, Vibratome, St. Lois, MO, USA) equipped with a sapphire blade in an ice-cold carboxygenated cutting solution containing (in mM): 110 choline, 25 NaHCO_3_, 3 KCl, 7 MgSO_4_, 1.5 NaH_2_PO_4_, 10 d-Glucose, 0.5 CaCl_2_.

Slices were incubated for one hour in a carboxygenated artificial cerebrospinal fluid (ACSF) composed of (in mM): 124 NaCl, 25 NaHCO_3_, 3 KCl, 1 MgSO_4_, 1.5 NaH_2_PO_4_, 10 d-Glucose, and 1.5 CaCl_2_ at 32°C, then transferred to a recording chamber and perfused with 36°C ACSF at a rate of 1-2 mL/minute. Neurons identified using DIC optics were recorded in whole-cell patch clamp mode using 4-6 MΩ glass pipettes filled with an intracellular solution (in mM: 130 K-gluconate, 10 EGTA, 10 HEPES, 1 CaCl_2_, 1 MgCl_2_, and 2 ATP, at pH 7.2-7.3; osmolality = 290-295 mOsm). Signals were amplified with an A-M Systems Model 2400 amplifier (Carlsborg, WA). Experimental protocols were conducted with pClamp software (v. 10.7; Molecular Devices, Sunnyvale, CA). An initial seal of greater than 1 GΩ, membrane resistance of greater than 100 MΩ, and positive action potential overshoot were inclusion criteria for seal and cell viability.

### Neuron identification

We visualized rNST under DIC optics with a Nikon E600FN microscope. EYFP was evident under epifluorescence but not adequate for targeting GABAergic neurons because it was expressed mainly in membranes and soma were obscured by dense neuropil labeling. Thus, neurons were classified based on their physiological response to light, i.e. optotagging (34–36), and as we have done in previous studies (12, 19). A fiber optic probe (200 μM, NA=0.39) connected to an LED (Thor Labs, model# 4100 4-channel LED Driver; 455 nm, 2.3mW at the tip) was centered to illuminate the rNST **(Fig. 1 inset**). Neurons were optically stimulated with 1 s light trains (10 Hz, pulse duration = 10 ms). Neurons were classified as non-GABAergic (G-) if they responded with long-latency IPSPs (> 9 ms) (see (19) for details). These IPSPs likely arose from directly-activated GABA neurons or inhibitory terminals in the nucleus. Two GABA-positive (G+) neurons were used in the dynamic clamp experiments (described below) and were identified by short-latency excitatory responses (< 3 ms). Photomicrographs of the patched neuron and its location were taken of the slice at high (40x) and low (4x) magnification under DIC and epifluorescence to aid in the localization of the neuron within the rNST.

**Fig 1.**
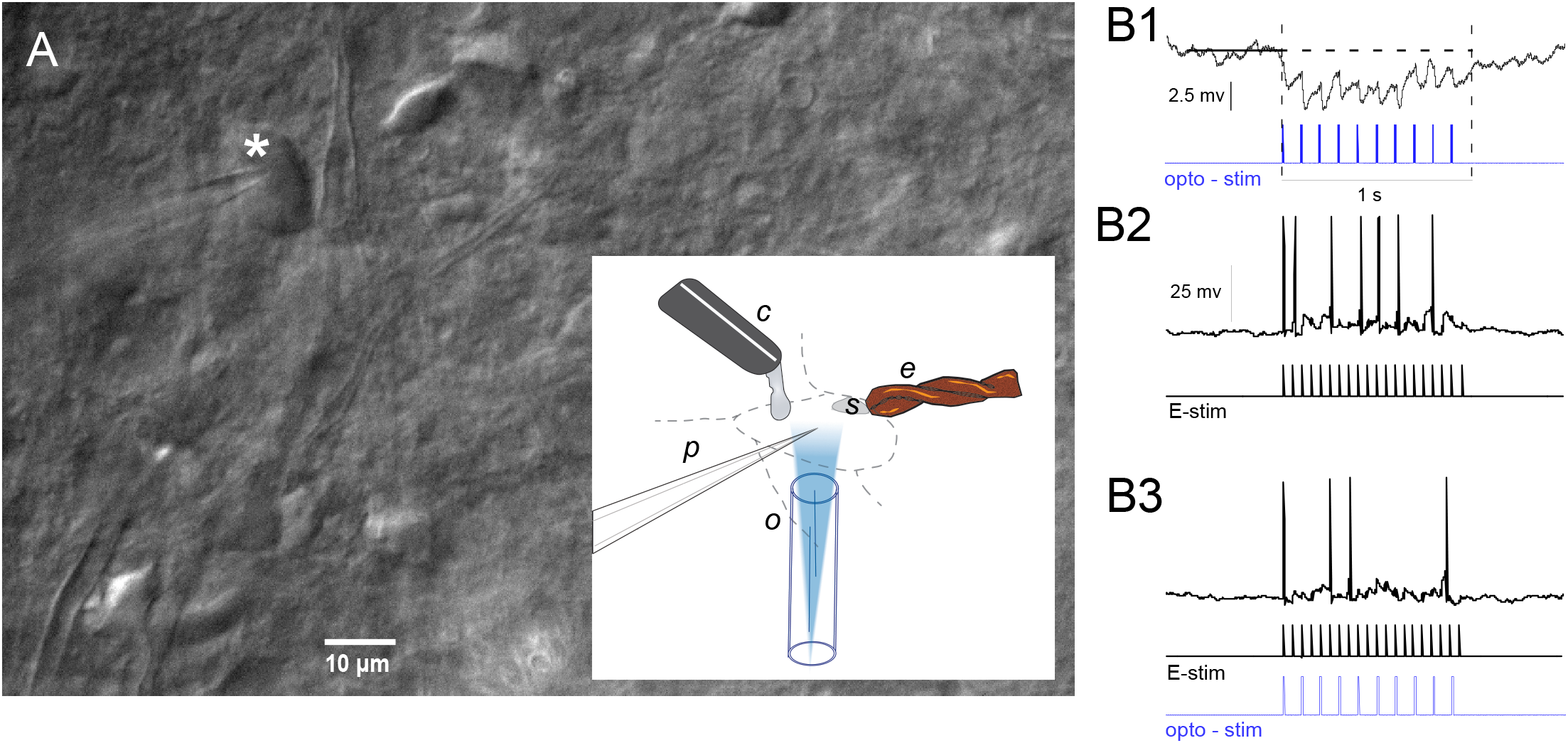
A. Photomicrograph of patched GABAergic negative cell under DIC optics (asterisk). Inset shows configuration for stimulation and recording through a pipette (p). A twisted bipolar stimulating electrode (e) was positioned over incoming afferent fibers in the solitary tract (s). A fiber optic probe was used for optogenetic stimulation (o) and a cannula positioned for drug delivery (c). B. Cells were categorized as non-GABAergic (putative projection cells) if they responded with inhibitory post-synaptic potentials (B1). The magnitude of the hyperpolarization was measured by subtracting the mean voltage level preceding optical stimulation from the mean level during optical stimulation. Action potentials were recorded in response to afferent fiber stimulation alone (B2) or coincident with optogenetic stimulation (B3).

### Current clamp

The presence of I_A_ in a recorded cell was identified in current clamp. The criterion was the occurrence of a delay in the latency of the first action potential, or an increase in the first ISI in response to a hyperpolarizing pre-pulse (450 ms duration, −100 pA to 0 pA in 20 pA steps) followed by a depolarizing pulse (1 s duration, 100 pA) (11, 12, 37, 38). Data presented in the current report include 7 I_A+_ and 5 I_A−_ cells from in an earlier paper (19), but incorporates new analyses including quantification of inhibitory postsynaptic potentials (IPSPs). Compound IPSPs were quantified under current clamp by subtracting the mean potential during a baseline period (0.5 s prior to stimulation) from that observed under optogenetic stimulation (see **Fig. 1B1**). This procedure did not measure the amplitude of individual IPSPs but captured the overall efficacy of GABAergic inhibition.

Responses of rNST neurons to afferent stimulation were quantified as the number of action potentials (APs) evoked by electrical stimulation of the solitary tract (ST; 150 μA,10 Hz,1 s, **Fig. 1B3**). The effect of GABA release on this response was determined by simultaneous optical stimulation (2.3 mW, 10 Hz, 10 ms duration pulses) in which trials with light stimulation were interdigitated with those without light stimulation (**Fig. 1B3**). Responses to GABA release were calculated as either absolute suppression (spikes control-spikes during light) or percent suppression (spikes control-spikes during light/spikes control * 100). The protocol was repeated whenever possible.

To determine the impact of I_A_ on responses to afferent stimulation and the modulation of these responses by optogenetic activation of the rNST GABA network, we recorded responses before and after adding the highly specific Kv4 channel blocker AmmTX3 (Alomone labs, STA-305) (32, 33) to the bath for a subset of neurons. AmmTX3 was previously shown to be highly effective in blocking I_A_ in rNST neurons (12). AmmTX3 (0.5 μM) was delivered at a rate of 0.5 ml/min through a 33 gauge stainless steel tube positioned ~300 μm from the dorsal edge of the rNTS (**Fig. 1A Inset**) and connected via PE 100 tubing to a 5 CC syringe in a syringe pump (Instech model 2000).

### Dynamic clamp

In a subset of experiments, dynamic clamp was used to determine if adding I_A_ to a cell lacking I_A_ (or after treatment with AmmTX3) could modulate the response to afferent stimulation. To implement the dynamic clamp protocol, we used values for I_A_ from a previous study that characterized I_A_ currents in rNST neurons under voltage clamp that employed both pharmacological and subtractive procedures to isolate the current (see (12)). From these recordings we obtained mean values of maximum I_A_ conductance (g_A_), and half-max parameters for activation (*θ_a_*) and inactivation (*θ_b_*), as well as inactivation decay (*τ_b_*) for G+ and G− neurons (see (12). These values were used to compute a synthetic I_A_ for real-time dynamic clamp.

The conductance-based model for I_A_ is given by *I_A_ = g_A_ a^3^ b (V_M_ − E_K_)*, where the gating variables *a* and *b* satisfy differential equations *a’ = (a_∞_(V_M_) − a) /τ_a_* and *b’ = (b_∞_(V_M_) − b) /τ_b_*, respectively. Here, 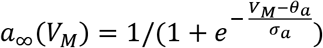 and 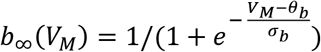. Parameter values for G+ (G−) cells are: *g_A_* = 9 10^−9^(1.2 10^−8^) *s*/*cm*^2^; *θ_a_* = −44(−52) *mV*; *θ_b_* = −69(−79) *mV*; *σ_a_* = 25.7(22.3); *σ_b_* = −1(−3); *τ_b_* = 140(136) *mS*; *τ_a_* = 2(2) *mS*; *E_K_* = −100(−100) *mV*.

As described at length in a previous publication, these parameter values recapitulated differences between G+ and G− neurons in the onset to AP initiation under current clamp (12).

We implemented dynamic clamp for I_A_ using the Linux-based Real-Time eXperimental Interface (RTXI v2.0) operating within Ubuntu 16.04.6 LTS (Xenial Xerus) on an Intel-Core i5-2500CPU @3.30Ghz x 4 processor equipped with an AMD CEDAR graphics card and National Instruments PCI-6221 Multifunction I/O card connected to the external input of the AM Systems Amplifier. Real-time computations occurred approximately every 0.25 microseconds, with peak computations taking no longer than 2.074 microseconds (RT Benchmarks Module, RTXI). Further details are provided in the Results. Software modules are available on Github (https://github.com/Trevor372/RTXI-DYNAMIC-IA).

### Statistical analysis

Paired t-tests were used to evaluate differences before and after a single treatment such as GABA releaser or the introduction of a synthetic I_A_ under dynamic clamp. A repeated-measures ANOVA with was appropriate when there were multiple treatments such as the impact of GABA infusion before and after infusion of AmmTX3. A significance level was set at P = .05 (Systat v. 13). Exact P values are reported in the Figure Captions or the Results text for values greater than .001. Error bars in the figures are standard errors of the mean (SEMs).

### Immunohistochemistry

At the conclusion of the experiment, tissue sections were fixed in 4% paraformaldehyde overnight then transferred to phosphate-buffered saline (PBS: 0.1 M, pH 7.4) the next morning. Slices were immunostained for the ionotropic purinergic receptor P2X2 to delineate the recording site relative to the terminal field of primary afferent fibers (39, 40). PBS rinses separated the following steps: Slices were treated with 1% sodium borohydride (20 min), then incubated in blocking serum (60 min, 0.3% Triton, 1% Bovine Serum Albumin, 5% Donkey Serum) prior to primary antibody incubation with a polyclonal rabbit P2X2 antibody (1:10000, Alomone Labs, Catalog#: APR-003, Lot#: AN-1502), followed by a fluorescently tagged anti-rabbit secondary antibody (Alexa Fluor 546, Catalog #: A10040, Lot#: 1833519). Photomicrographs were taken with a DS-Ri1 Nikon camera attached to a Nikon Eclipse 600 microscope using brightfield and fluorescent illumination with appropriate fluorescent filters. These images were compared to the photos taken during recording. Locations of rNST cells were plotted on a representative section midway between the rostral pole of the nucleus and the level at which the nucleus is no longer adjacent to the 4^th^ ventricle. Locations were approximated by comparing the location of the recording electrode to the P2X2 field and other landmarks including the borders of the nucleus, the ST, and myelinated fibers coursing through the middle of the section.

## Results

### Afferent-induced responses were preferentially suppressed by GABA in I_A_-positive neurons

We tested the effects of the optogenetic release of GABA on the response to afferent stimulation in 57 G− neurons (e.g. **Fig. 1B2** and **1B3**). Although the response to afferent stimulation was reduced both in cells with and without I_A_ (paired t-tests: I_A_+, P < .001, N =30; I_A_−, P <.001, N = 27), the percent suppression was greater in G− cells with I_A_ (I_A_+: 62.1 ± 5.6%; I_A_−: 38.8 ± 4.9%; t-test: P = .003, N: I_A_+ = 30, I_A_− = 27; **Figs. 2A**).

**Fig. 2.**
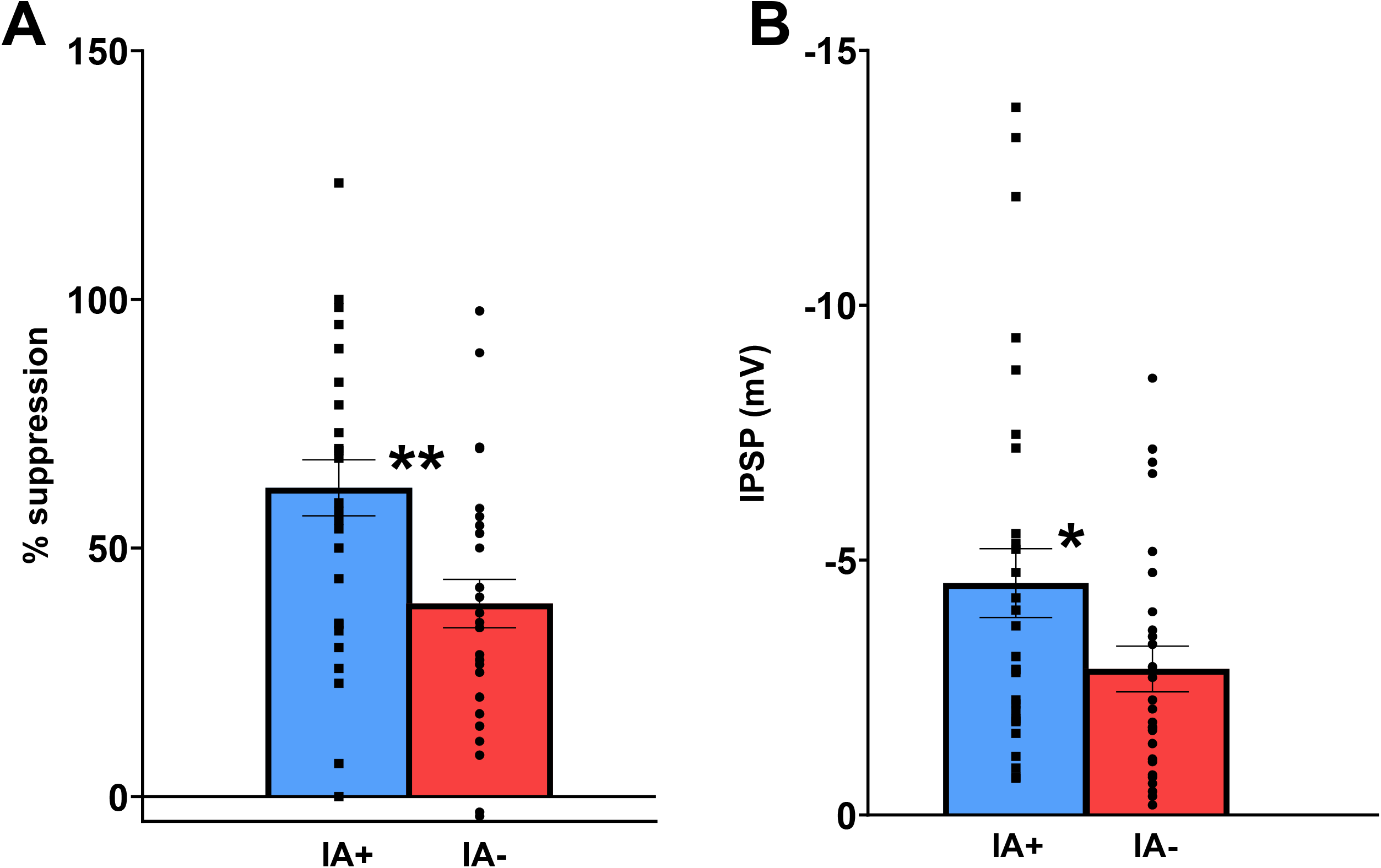
A. The percent suppression in the number of APs following afferent stimulation associated with optogenetic release of GABA was greater in cells with I_A_ compared to those without (** t-test: P = .004, N: I_A_+ = 30; IA− = 27). B. The mean amplitude of the compound IPSP following optogenetic release of GABA was greater in cells with I_A_ compared to those without (* t-test, P = .039, N: I_A_+ = 30, I_A_− = 27).

Across all G− neurons, the size of the compound IPSP following optogenetic release of GABA correlated positively with the degree of suppression in the response to afferent stimulation (R = 0.49, P < .001) and the mean amplitude of the compound IPSP greater in cells with I_A_ (t –test, P = .043) (**Fig. 2B**). This supports the hypothesis that one cause of the differential suppression in I_A_+ and I_A_− neurons is the size of the optogenetically-induced IPSP. However, as evident from **figure 2B**, there was considerable variability in IPSP magnitude, suggesting that additional factors may contribute. For example, I_A_ could be contributing to the enhanced inhibition observed in I_A_+ G− neurons by a feed-forward mechanism. In such a scenario, feed-forward GABA-mediated hyperpolarization would further de-inactivate I_A_ channels in the projection cell such that the excitatory afferent input would be blunted by the activated outward I_A_ current. To explore this potential interaction, we compared the response to afferent stimulation under control conditions and during blockade of I_A_ channels with AmmTx3 and the degree to which the response to afferent stimulation was suppressed by optogenetic activation of NST GABA.

### Pharmacological block of I_A_ increases the response to afferent stimulation

Of the 57 cells in which we assessed the effects of optogenetically released GABA, 25 (17 I_A_+; 8 I_A_−) were tested further following infusion of AmmTX3 into the slice chamber (**Figs. 3A1 & 3A3**). In neurons with I_A_, AmmTX3 produced a pronounced increase in the response to afferent stimulation (53.27%, paired t-test: P = .001, **Fig. 3B left**) but did not affect the afferent response in cells without I_A_ (−7.49%. NS, N = 8, **Fig. 3B right**). We further observed that I_A_+ cells were more responsive to afferent stimulation (6.77 ± 1.27 spikes/s) compared to cells without (3.28 ± 0.78 spikes/s, t-test, P = .028). This is consistent with our previous study that showed a trend in the same direction (Chen, 2018). However, in that study, with a smaller N, the effect did not reach significance.

**Fig. 3.**
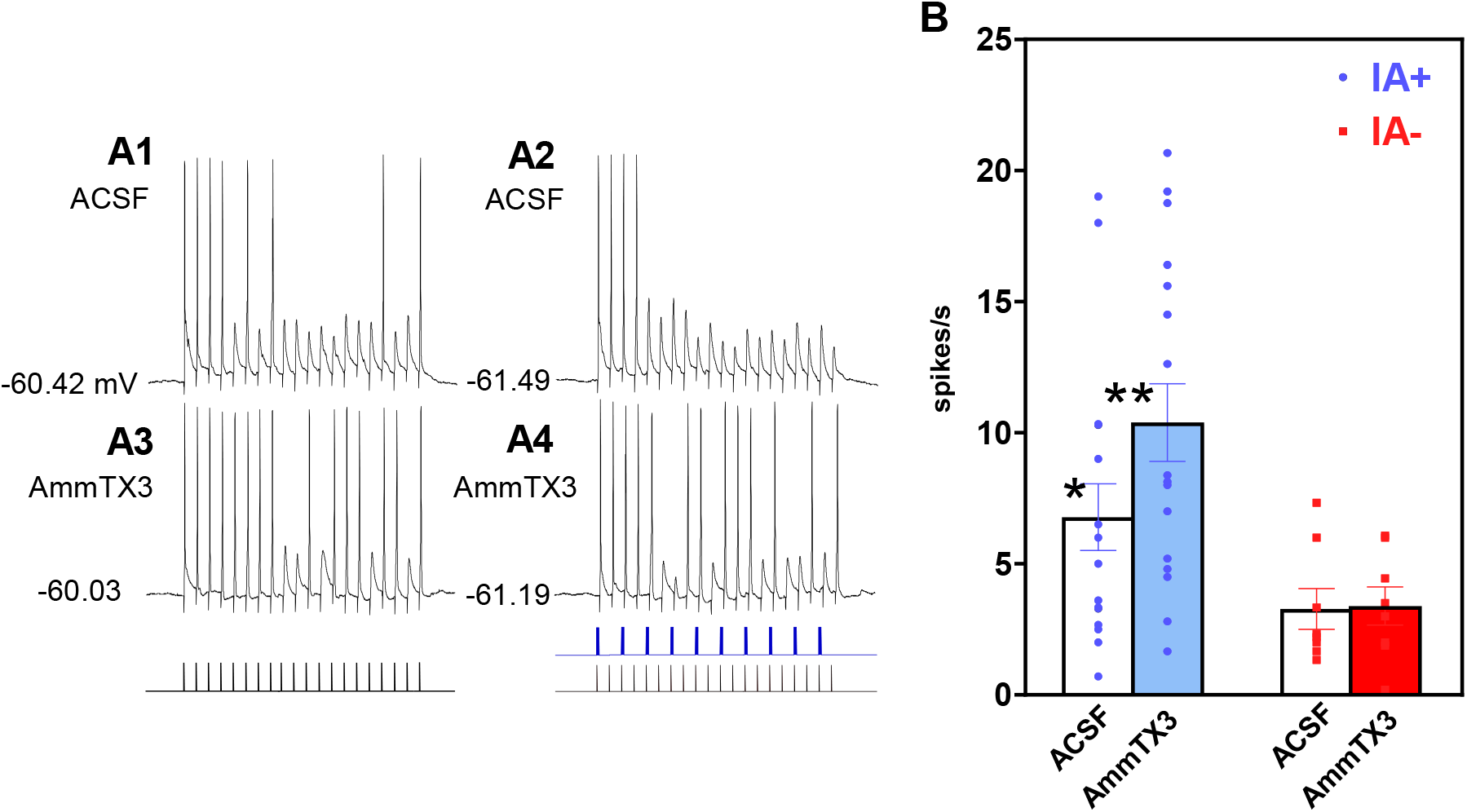
APs elicited by afferent stimulation in a cell with I_A_ (A1) were reduced following optogenetic release of GABA (A2) but increased following drug treatment with AmmTX3 (A3) and when combined with optogenetic stimulation (A4). B. The mean number of APs elicited by afferent stimulation increased by 53% after drug treatment with AmmTX3 in cells with I_A_ (** t-test, P = .001) but were unchanged in those without (t-test, P > .05). In addition, under control conditions (ACSF), the response to afferent stimulation was greater in cells with I_A_ compared to those without (* t-test, P = .028, N: I_A_+ = 17, I_A_− = 8).

This increase in excitability to afferent stimulation in I_A_+ cells after AmmTX3 could not be explained by alterations in the resting membrane potential (RMP) which did not change in the two states (control: − 58.59 ±1.51 mV; AmmTX3: −56.18 ± 1.68 mV; P = .18, N = 17). Nor was there a significant correlation between the difference in RMP before and after drug treatment and the change in spike count (R = 0.069, P = .548). It was notable, however, that there were significant changes in action waveform following drug treatment (**Table 1**), including a widening of the AP half-width and a reduction in the afterhyperpolarization (AHP).

**Table 1.** Action Potential Waveform Before and After AmmTX3

|  | Control | AmmTX3 | P |
| --- | --- | --- | --- |
| RMP (mV) | -58.59 (1.51) | -56.18 (1.68) | 0.18 N = 17 |
| Peak (mV) | 71.68 (2.8) | 70.69 (3.91) | 0.744 N = 12 |
| AHP (mV) | -18.87 (1.97) | -14.43 (1.58) | 0.008 N = 12 |
| Half-width (ms) | 1.13 (.095) | 1.33 (0.11) | 0.008 N = 12 |
| MR (MΩ) | 442.2 (43.68) | 411.6 (51.74) | 0.366 N = 24 |

Values are means (SEM). RMP, resting membrane potential, AHP, after hyperpolarization, MR, membrane resistance.

### Injecting synthetic I_A_ with dynamic clamp reduces excitability to afferent stimulation

To further assess a causal role for I_A_ in modulating rNST responses to afferent stimulation, we implemented dynamic clamp to induce an I_A_ conductance (**Fig. 4A**). The parameters for this synthetic I_A_ were based on previous empirical studies in which we characterized I_A_ kinetics and then showed that these parameters reproduced a delay to spike under dynamic clamp using a protocol of depolarization preceded by hyperpolarization (see Fig. 11, Chen, 2020). In 11 cells that either lacked constitutive I_A_ (N = 6) or I_A_+ cells in which I_A_ was blocked pharmacologically (N = 5), a synthetic I_A_ current was inserted into cells during afferent stimulation. On average, synthetic I_A_ reduced the response to 20 Hz afferent stimulation by about 14% (6.96 ± 1.68 vs 6.06 ± 1.65 spikes/s; t-test: P = .004). Indeed, in all but one cell, a dynamic clamp I_A_ conductance reduced the number of spikes in response to afferent stimulation **(Fig. 4B)**. Thus, although the magnitude of the effect was small it was highly reproducible **Fig. 4C**).

**Fig. 4.**
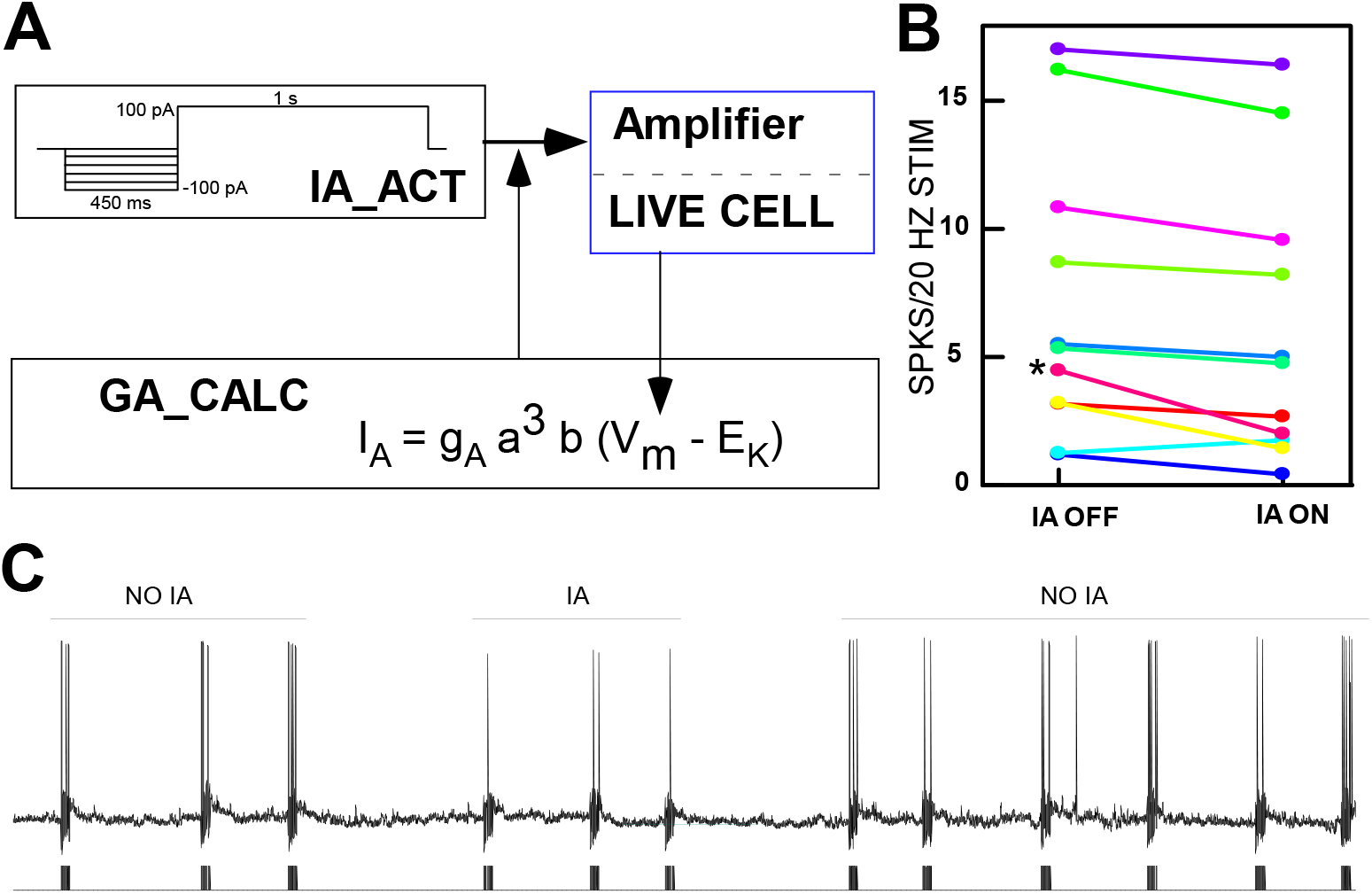
A. Schematic of the method for implementing dynamic clamp. Membrane voltage from cells either lacking constitutive I_A_ or I_A_ blocked with AmmTX3 were used to compute a synthetic IA conductance employing parameters for I_A_ derived from empirical studies (12). B. Introducing I_A_ suppressed responses to afferent stimulation in 10/11 cells. The effect was small but consistent. An example from the cell with the most marked effect (asterisk in B) is shown in (C).

### Impact of AmmTx3 on GABAergic suppression

The activation of I_A_ channels suppresses afferent-evoked responses but does activation of I_A_ channels interact with GABA-induced inhibition, which also functions to suppress afferent input? If I_A_ is causal to the enhanced GABA inhibition in I_A+_ cells, blocking this current would result in a loss of GABA efficacy. However, this did not appear to be the case. Optogenetic release of GABA produced a significant reduction in the response to afferent stimulation in I_A_+ cells both before and after treatment with AmmTx3 (**Fig. 3A2 & 3A4, Fig. 5A**) (ANOVA: AmmTX3, P <.001; GABA, P < .001, AmmTX3 x GABA, P = .372, N = 17). Moreover, although there was an apparent decrease in the % suppression after blocking I_A_ (paired t-test, P = .025, N = 17, **Fig. 5B1**), this effect was dependent on two cells in which the response to afferent stimulation was entirely eliminated by GABA activation prior to AmmTX3, making the extent of the actual suppression unknown. When these cells were removed from the analysis, the effect was lost (P = .069, N = 15). Indeed, the apparent decrease in GABA efficacy following treatment with AmmTX3 was likely due to the increased excitability of cells to afferent stimulation rather than a decrease in GABA suppression. When the magnitude of optogenetic inhibition was calculated as the difference in the absolute number of spikes (**Fig. 5B2**, Δ spikes), AmmTX3 had no effect, i.e. blocking I_A_ did not change the efficacy of GABA to reduce cell spiking.

**Fig. 5.**
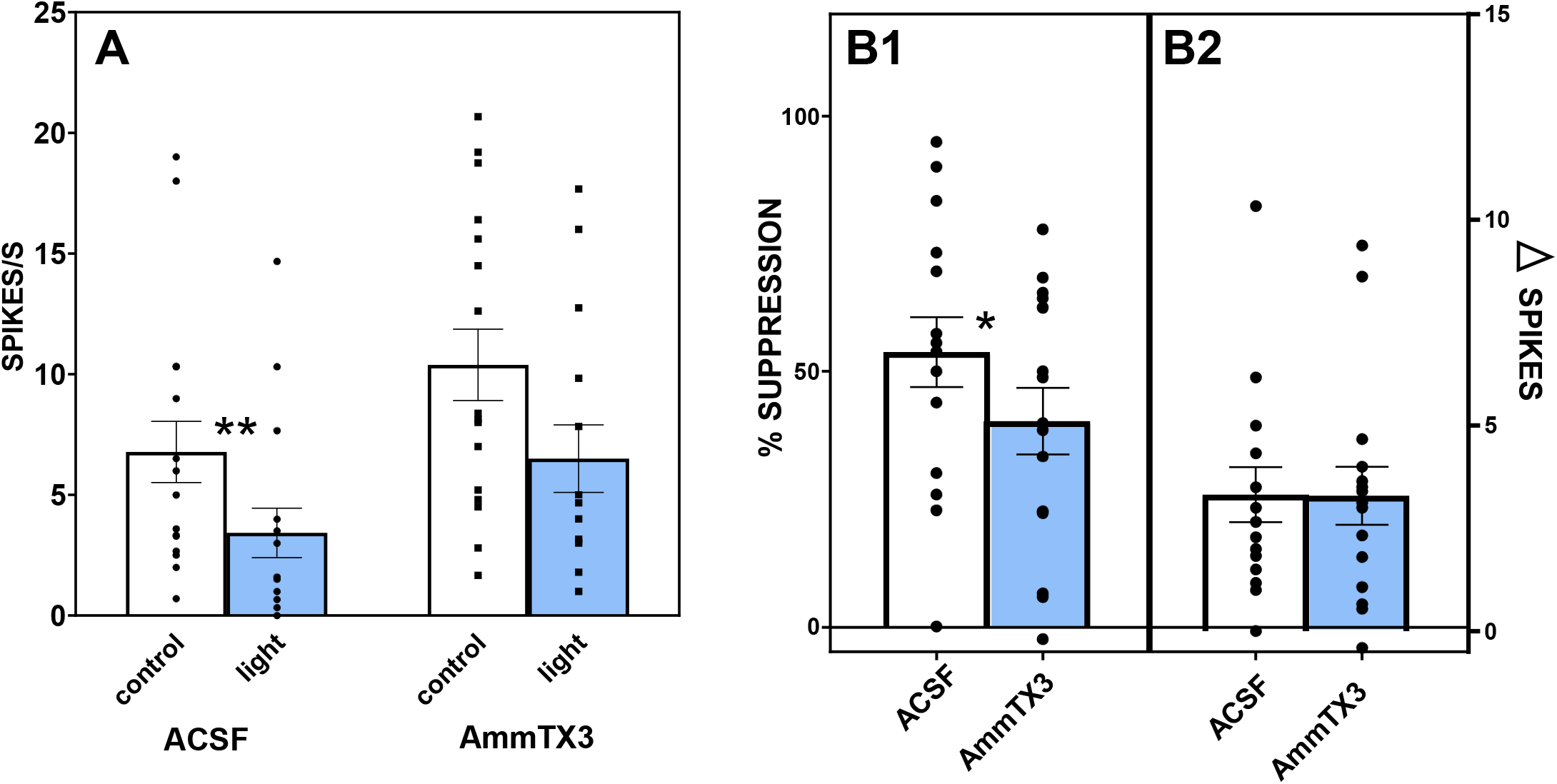
A. Following optogenetic release of GABA (light) there was a significant reduction in the ST-evoked response compared to control in IA+ cells both before and after treatment with AmmTX3. (** ANOVA: drug, P <.001; GABA, P < .001, drug x GABA, P = .372, N = 17). B1. Despite the lack of a significant interaction between GABA and AmmTX3 using spike counts, the mean percent suppression due to GABA was smaller following AmmTX3 (paired t-test, P = .025, N = 17). However, this only approached significance when 2 cells with a floor effect (prior to drug treatment) were removed (* P = .069, N = 15). B2. Moreover, when the magnitude of inhibition was calculated as the difference in the absolute number of spikes before and after optogenetic stimulation (spikes), AmmTX3 had no effect.

### Location of cells responding to afferent stimulation

Many of the cells with I_A_ responding to afferent stimulation were in the central subdivision (10/25). Three of the 8 in the medial subdivision expressed I_A_, 2/5 in the ventral subdivision, and 2/2 in the lateral subdivision (**Fig. 6**). Many of the cells outside the central subdivision tended to be located adjacent to the central subdivision defined by the presence of P2X2 (when available).

**Fig. 6.**
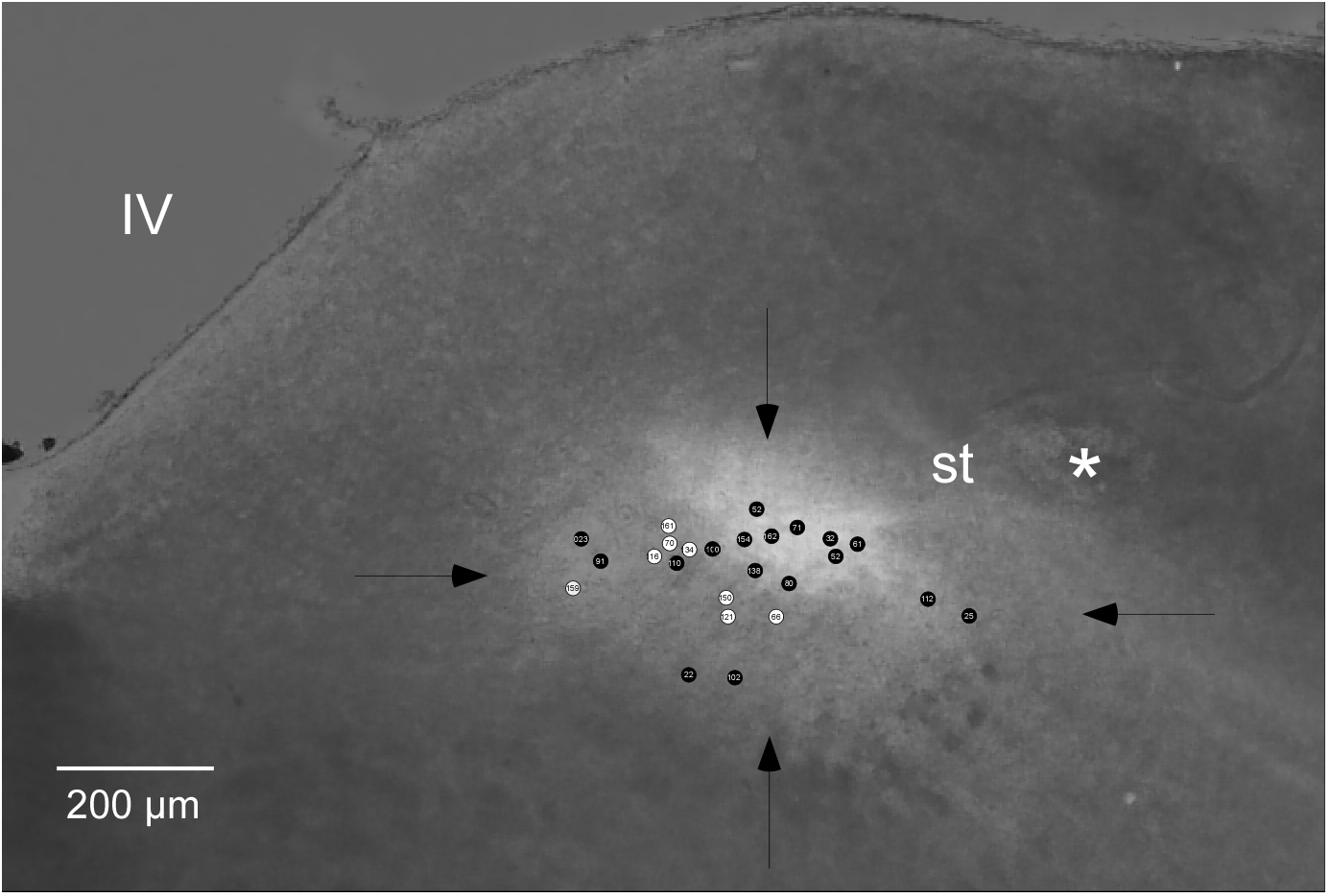
Photomicrograph of the rostral solitary nucleus stained for P2X2 which effectively demarcates the central subdivision. Approximate borders of the nucleus are demarcated with arrows. Incoming fibers of the solitary tract (st) were also labeled with P2X2. Many cells with I_A_ were located in the central subdivision (black symbols). Despite the failure to identify cells in the central subdivision without I_A_ in this sample (white symbols), such cells were recorded in our previous study (12). An asterisk designates damage to the tissue where a stimulating electrode was positioned adjacent to the solitary tract. IV, 4th ventricle.

## Discussion

The results of the present study suggest that GABAergic inhibitory synapses and the transient, rapidly inactivating outward K^+^ current (I_A_) function independently to regulate the purely excitatory input of gustatory afferent fibers into the rNST. Compound IPSPs evoked by the optogenetic release of GABA were larger in cells with I_A_ compared to those without, and thus provide a partial explanation for the greater suppressive effect of GABA on rNST responses in cells with I_A_. Although there is evidence for GABAergic feed-forward inhibition in the rNST, this inhibition does not appear to potentiate an I_A_ current activated by excitatory afferent input. Specifically, blocking Kv4 channels with AmmTX3 did not modulate GABAergic inhibition. However, blocking Kv4 channels greatly enhanced the response to afferent stimulation, suggesting the presence of a window current. Thus, I_A_ can suppress excitatory input even in the absence of hyperpolarization afforded by inhibitory input. These results were corroborated using dynamic clamp to demonstrate that adding a synthetic I_A_ suppressed the response to afferent stimulation.

### GABA and I_A_ in the rNST

The presence of both GABA and I_A_ have been well characterized in the rNST. Immunohistochemical visualization of GABA delineates the outline of the nucleus and GABA interneurons are estimated to account for as many as 36% of rNST neurons (41). In addition, GABA from extrinsic sources includes the amygdala and more caudal, viscerally-related NST (16, 18, 42). GABAergic synapses are implicated in gain control, the tuning of gustatory profiles, and may contribute to metabolic modulation of taste responses (10, 23, 43, 44). For example, when GABA_A_ receptors were blocked with bicuculline, neurons became more broadly responsive to gustatory stimuli (23), suggesting that afferent fibers carrying sideband sensitivities were under tonic inhibition. Similarly, optogenetic activation of GABAergic neurons in the rNST differentially suppressed sideband sensitivity compared to the best stimulus (24). Other studies have shown that blocking GABA_A_ receptors increases the magnitude of excitatory responses (29, 45), suggesting a role in gain control.

I_A_ is also well represented in the rNST, and pharmacological block with either 4-AP or the specific KV4 channel blocker AmmTX3 suppressed I_A_ currents (11, 19). Indeed, I_A_ currents have been identified in both rNST projection neurons and GABAergic interneurons, although a subset of GABAergic neurons in the ventral subdivision appear not to express this current (25, 26, 46). Despite the robust presence of I_A_ in the rNST, a functional role is unclear. In the cNST, I_A_ is more pronounced in neurons with projections to the hypothalamus compared to those with local brainstem projections to the ventrolateral reticular formation (28, 30). Because local projection cells more faithfully follow afferent nerve stimulation compared to those with ascending projections, it has been proposed that I_A_ in cells with ascending projections may provide a mechanism for state-dependent modulation of visceral signals with a hypothalamic destination, compared to neurons with a purely reflex function (28). Indeed, the expression of channels supporting I_A_ are subject to metabolic state (47–49). However, the distinction between cells with local brainstem projections compared to those with ascending projections is not as pronounced in the rNST. Approximately 72% of PBN projection cells express I_A_ compared to 58% of cells projecting to neurons involved with reflex-like consummatory ingestive responses in the subjacent RF (25, 26). Both populations of projection cells receive monosynaptic afferent input from fibers in the VIIth and IXth cranial nerves (25, 26). Because this input is glutamatergic and involves convergence of multiple afferent fibers onto second-order neurons (22, 29), I_A_ channels may be an important mechanism for limiting excess excitability.

### Regulation of excitatory input by IA

Hoffman and colleagues showed a gradient of I_A_ in hippocampal pyramidal neuron dendrites that increased from proximal to distal (27). Blocking I_A_ with 4-AP increased the amplitude of an evoked excitatory postsynaptic potential (EPSP) by over 50%, suggesting that I_A_ channels acted as a “shock absorber” to regulate excitatory inputs. Although we were unable to measure EPSP amplitudes following treatment with AmmTX3 due to the presence of APs, we note that the increase in excitability reflected in AP frequency was just over 50%. Similarly, Cai and colleagues showed that focal application of 4-AP at the branch point of terminal dendrites of hippocampal pyramidal cell facilitated AP firing (50).

In addition to suppressing dendritic EPSPs, I_A_ channels may influence excitability by altering AP waveform (11, 51–55). Similar to a previous report using 4-AP to block I_A_ channels in rNST neurons (11), the specific I_A_ channel blocker AmmTX3 produced an increase in the AP half-width, a decrease in the AHP with no change in the resting membrane potential. However, despite these changes in spike waveform the results of our previous rNST study did not find a change in the AP threshold to injected current, nor an increase in the number of APs elicited by injected current following treatment with AmmTX3 (12). Thus, it seems likely that the increase in the responsiveness to afferent stimulation observed in the current study after blocking KV4 channels likely reflects a role for I_A_ in dendritic excitability that ultimately influences AP initiation.

An I_A_ window current, i.e., one that is operational at the resting membrane potential (56–58), is evident in both the cNST and rNST by the overlap of activation and inactivation curves for isolated I_A_ currents (12, 30, 31). Evidence that this mechanism can directly modulate excitability is the observation of a shortened latency to AP initiation in response to a depolarizing stimulus in the absence of a hyperpolarizing pre-pulse (12). In the present study, the increase in responsiveness to afferent stimulation following treatment with AmmTX3 further suggests that an I_A_ current was active at rest.

### Interaction between GABA and I_A_

Bradley and colleagues were the first to propose a feedforward circuit in the rNST based on the presence of IPSPs in 2^nd^ order rNST neurons following afferent stimulation, as well as enhanced EPSPs when GABAergic inhibition was blocked (6, 11, 29). In addition, there is evidence that rNST GABAergic neurons receive monosynaptic afferent input (41, 59). Thus, with a window current close to the RMP, a phasic inhibitory input might be expected to further de-inactivate I_A_ channels such that incoming excitatory afferent input that opens the outward K+ channel would add to the suppressive effect of the phasic inhibition itself (30, 60). Despite the evidence for a feed-forward inhibitory circuit and a widespread substrate for both GABA inhibition and I_A_ currents in rNST, we saw little evidence for an interaction between these two regulatory mechanisms. When we factored out the general increase in excitability when I_A_ was blocked, and calculated only the absolute number of spikes associated with inhibition, there was no evidence for decreased inhibition in the absence of functioning KV4 channels.

The lack of a dynamic interaction between I_A_ and GABAergic inhibition in the rNST may reflect the location of these respective channels and receptors. I_A_ channels in the hippocampus and cortex are located in close proximity to GABAergic synapses (61, 62) and an interaction between these two mechanisms in the cortex was shown by blocking I_A_ in cortical pyramidal cells with 4-AP (63). This not only increased the amplitude of a back-propagated calcium signal in proximal dendrites, but also increased the magnitude of a GABA-induced IPSP. Likewise, an interaction between GABAergic inhibition and I_A_ was observed in midbrain dopaminergic neurons (64), which have inhibitory inputs to both dendritic and somal compartments (65). In this region, the pharmacological block of I_A_ currents with AmmTX3 shortened delays associated with high-frequency stimulation (postburst delays) that were then entirely suppressed by blocking GABA, i.e. both mechanisms contributed to postburst delays. We did not observe a change in the amplitude of the IPSP to optogenetic release of GABA following AmmTX3, nor did we see a decrease in the efficacy of GABAergic inhibition to suppress response to afferent stimulation. However, in the rNST, KV4.3 channels predominate in the neuropil of the rNST (12) and may be concentrated on distal dendrites as seen in hippocampal pyramidal neurons (27, 66). Although distal dendrites are also the location of excitatory afferent inputs to the rNST that might activate KV4 channels, they are distant from GABAergic synapses that are more dominant on soma and more proximal dendrites (67), making interactions less likely. Nevertheless, despite the lack of a dynamic interaction between GABAergic inhibition and I_A_, it seems clear that an I_A_ window current works in concert with GABAergic feed-forward and tonic inhibition to regulate afferent responses in the rNST.

Although our evidence to date suggests that I_A_ and GABAergic pathways work independently to regulate rNST responses to afferent stimulation, this conclusion is subject to several methodological considerations. GABAergic synapses in the rNST derive from multiple sources including both intrinsic local and external sources. To this we might add excitatory inputs to the rNST that synapse on GABAergic interneurons reviewed in (68). Our optogenetic activation of channelrhodopsin in GABAergic neurons/terminals did not discriminate among these sources. As such, full-on release of GABA might have obscured more subtle interactions with I_A_ had only one source of GABA been activated. Additional studies focused on specific GABAergic pathways may clarify if there are any conditions under which GABA mediated hyperpolarization interacts with I_A_.

## Acknowledgments

The Venus fluorescent protein was developed by Dr. Atsushi Miyawaki at the RIKEN Brain Science Institute in Wako Japan; the VGAT-VENUS transgenic mouse developed by Dr. Yuchio Yanagawa was made available to us from Dr. Hisashi Umemori at the University of Michigan Medical School, Ann Arbor Michigan. Excellent technical help was provided by Andrew Harley and Trevor Haas. Thanks also to Charlotte Klimovich for her artistic contribution to the figures and for excellent editorial assistance. We are also greatly indebted to Dr. Sharmila Venugopal for her thoughtful insights and suggestions.

